# Cellular and Cytokine Responses in the Granulomas of Asymptomatic Cattle naturally infected with *Mycobacterium bovis* in Ethiopia

**DOI:** 10.1101/2020.06.12.149518

**Authors:** Begna Tulu, Henny M Martineau, Aboma Zewude, Fekadu Desta, David A Jolliffe, Markos Abebe, Taye Tolera, Mulugeta Belay, Adrian R Martineau, Gobena Ameni

## Abstract

Cellular (CD3+ T cell and CD68+ macrophages), cytokines (IFN-γ+ and TNF-α+) and effector molecule (iNOS+) responses were evaluated in the lymph nodes and tissue of cattle naturally infected with *Mycobacterium bovis*. Detailed post mortem and immunonohistochemical examinations of lesions were performed on 16 cows positive for single intradermal cervical comparative tuberculin test (SICCTT) and identified from dairy farms located around the Addis Ababa City. The severity of the gross lesion was significantly higher (p=0.003) in *M. bovis* culture positive (n=12) cows than in culture negative (n=4). Immunohistochemical techniques showed that the mean percentage labelling of CD3+ T cells decreased as the stage of granuloma increased from stage I to stage IV in culture positive cows (p<0.001). On the other hand, the proportional fraction of CD68+ macrophages and the concentrations of IFN-γ+, TNF-α+ and iNOS+ increased significantly from stage I to stage IV (p< 0.001) in culture positive cows. At the early stage of the granuloma, the culture negative cows showed significantly higher mean proportions of CD68+ macrophages (p=0.03) as well as the concentrations of iNOS+ (p=0.007) compared to culture positive cows. Similarly, at advanced granuloma stages, culture negative cows demonstrated significantly higher mean proportions of CD3+ T cells (p< 0.001) compared to culture positive cows. Thus, the present study demonstrated that following natural infection of cows with *M. bovis*, as the stage of granuloma increases from stage I to stage IV, the proportion of CD3+ cells decreases while the immunolabeling fraction of CD68+ macrophages, IFN-γ+, TNF-α and iNOS+ increases.

## INTRODUCTION

Bovine tuberculosis (bTB) is a chronic infectious disease of cattle mainly caused by *M. bovis*, a member of the *Mycobacterium tuberculosis* complex (MTBc). *M. bovis* has a wide host range that includes domestic animals, wildlife and humans (1, 2). With over 50 million infected cattle worldwide, bTB causes significant economic loss to the agricultural industry, costing US$3 billion annually (3). Effects on human morbidity and mortality are also considerable. In 2019 alone, it was reported that *M. bovis* was responsible for 143, 000 new human TB cases and 12, 300 deaths. Over 91.0% of the deaths were from African and Asian countries (4).

In some developed countries, the introduction of test and slaughter of bTB infected cattle together with continuous surveillance systems and movement restrictions, has achieved dramatic results in lowering the prevalence and even eradicating the disease (5, 6). However, these control programs are costly, and in countries like Ethiopia where bTB is an endemic disease and the agricultural economy relies on traditional farming practices (7, 8), new tools like effective vaccination and immunodiagnostic are urgently needed (2, 9, 10).

The Single intradermal cervical comparative tuberculin test (SICCTT) is the most widely used test for the diagnosis of bTB in live cattle (11). SICCTT measures the delayed hypersensitivity reaction to the tuberculin antigen-purified protein derivative (PPD) of *Mycobacterium bovis* (PPDb) and *Mycobacterium avian* (PPDa). In infected animals, there is swelling and indurations at both injection sites 72hr later (11, 12). However, SICCTT has lower sensitivity when there is co-infection with certain parasites like *Fasciola hepatica and Strongylus sp* (13, 14) which are widely distributed in Ethiopia (15, 16).

The second feasible bTB control option for developing countries like Ethiopia is through the vaccination program. However, presently, there are no effective vaccines that exist for the control of bTB in cattle. Bacillus Calmette Guerin (BCG) which is used in humans has certain limitations in cattle, including interference with SICCTT test.

Hence, understanding the local immunological responses is of paramount importance in the effort to develop new vaccines and diagnostic tools (2, 9). During mycobacteria infection, granuloma formation is the main mechanism of host immune response to contain the spread of bacterial dissemination, but this can result in significant tissue damage (17, 18). Immunity against mycobacteria is primarily a cell mediated immune (CMI) response, which involves recruitment of macrophages, dendritic cells, and helper T cell type-1 (TH1) modulated by cytokines (17, 19, 20). Cytokines like interferon gamma (IFN-γ) (20), interleukin-12 (IL-12) (21), IL-6, and tumor necrosis factor (TNF) play a significant role in activating immunological cells to kill mycobacteria and inducing TH-1 responses (22). In addition, the production of molecules like nitric oxide (NO) by macrophages or phagocytic cells during mycobacterial infection play a crucial role in the intracellular killing of mycobacteria as it is cytotoxic at high concentrations. NO release is enhanced by inflammatory stimuli via the up regulation of the inducible for of NOS (iNOS or NOS2) with in inflammatory macrophages (23, 24). Conversely, cytokines such as IL-4 (25) and IL-10 (26), known as the anti-inflammatory cytokines, are responsible for down-regulating the role of pro-inflammatory immune responses to control the tissue damage (17).

Existing studies on the immune response of cattle against *M. bovis*, largely focus on the experimental infections generated through the respiratory route (10, 17, 27-29). Through characterization of gross and microscopic lesion development, these studies have shown host immune response related factors to influence bTB diseases outcome (19, 30). Host genetic makeup and age related factors have also been shown to influence the susceptibility to *M. bovis* infection (31, 32).

However, there are few studies on such fundamental aspect of host immune response in a natural infection setup (33, 34). Menin *et al*., (2013) describe that during natural infection with bTB, the lesion severity, measured using a pathology severity score (33), is positively correlated with viable bacterial loads. Similarly, neutrophil numbers in the granuloma are associated with increased *M. bovis* proliferation (33). As the stage of granuloma increases, the expression of cytokines increases mediated by macrophages and epithelioid cells (35). More specifically, little is known about the local immune response of CD3+ T cells, CD68+ macrophages, IFN-γ, TNF-α and iNOS in cattle naturally infected with *M. bovis*. Thus, the objective of this study was to evaluate the responses of selected (CD3+ T cells and CD68+ macrophages) immune cells and pro-inflammatory cytokine (IFN-γ, TNF-α) and and effector molecule (iNOS) across stages of granuloma in cattle with natural *M. bovis* infection.

## RESULT

### Animals and their *M. bovis* culture status

A total of 16 cows with the mean age of 5.8 years (age range from 2.5 to 9 years) which were positive for SICCT test (≥ 2 mm cut value), were considered for the analysis. Regarding their body condition, 7 (44.0%) of the cows had poor, 6 (37.5%) medium and the rest 3 (18.7 %) had good body condition. Twelve (75.0%) of the cows were positive for *M. bovis* culture while 4 (25.0%) were negative for *M. bovis* culture (Table S1). bTB suggestive lesions were detected in 132 (48.5%) samples from which 99 (56.3%) were lymph nodes from the head and neck region, thorax, and abdomen and 33 (34.4%) were from lung tissues.

### Gross pathology

Based on the postmortem examination, all 16 cows had gross lesions in the following lymph nodes; caudal mediastinal 16 (100.0%), bronchial 15 (94.5%) and cranial mediastinal 13 (81.3%). Six (37.5%) cows also had lung lesions. The majority of the lesions were characterized by caseous necrosis. The median severity of the gross pathology was significantly greater (p=0.004) in *M. bovis* culture positive animals than in *M. bovis* culture negative animals (Fig. 1C). Within culture positive cows the severity of the lymph node lesions was significantly higher in the thoracic lymph nodes (p<0.05) as compared to head and abdominal lymph nodes (Fig. 1A).

**Figure 1:**
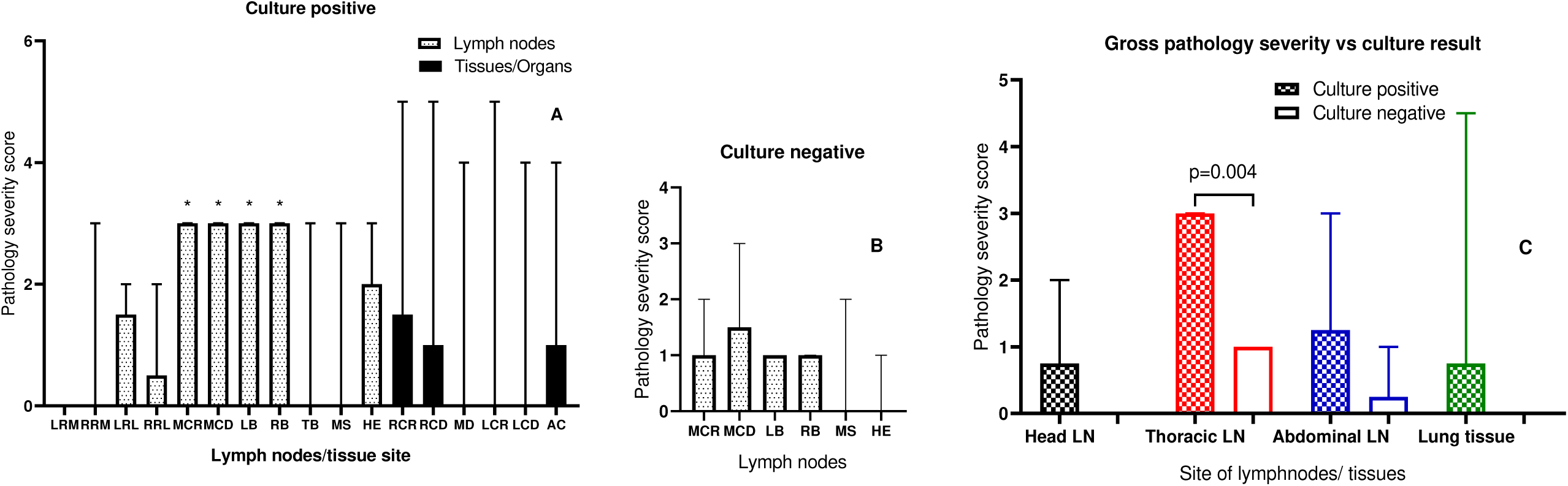
Gross pathology severity score of lymph nodes and lung tissues of cattle positive for *M. bovis* culture (n=12) compared to negative for *M. bovis* culture (n=4). A) Gross pathology severity score of the *M. bovis* culture positive animals. *p<0.05. B) Gross pathology severity score of the *M. bovis* culture negative animals. C) Gross pathology severity score vs. culture result of both culture positive and culture negative animals. Lymph nodes, LRM: left retropharyngeal medial, RRM: right retropharyngeal medial, LRL: left retropharyngeal lateral, RRL: right retropharyngeal lateral, MCR: medial cranial, MCD: medial caudal, LB: left bronchial, RB: right bronchial, TB: tracheobronchial, and MS: Mesenteric, HE: hepatic, and lung, RCR: right cranial, RCD: right caudal, MD: middle, LCR: left cranial, LCD: left caudal, and AC: accessary.

### Histopathology

A total of 37 tissues were examined from both culture positive and culture negative animals. Representative microscopic findings are shown below (Fig. 2). The four culture negative cows had granulomas in their cranial and caudal mediastinal lymph nodes only. In culture positive animals, a larger number of granulomas and diversified stages of granulomas were detected. Caudal and cranial mediastinal lymph nodes showed the majority of the histopathology results generated in this study (Table 1).

**Table 1:** Distribution of granulomas (stage I-IV) of selected lymph nodes and lung tissue of examined for 16 animals.

| Types of LN/Tissue examined | Total No. of granuloma (%) | Stages of granuloma |  |  |  |  |  |  |  |
| --- | --- | --- | --- | --- | --- | --- | --- | --- | --- |
|  |  | Culture positive |  |  |  | Culture negative |  |  |  |
|  |  | I | II | III | IV | I | II | III | IV |
| Retropharyngeal | 11 (13.0) | 2 | 4 | 3 | 2 | 0 | 0 | 0 | 0 |
| Cranial mediastinal | 15 (17.6) | 1 | 2 | 4 | 5 | 0 | 1 | 1 | 1 |
| Caudal mediastinal | 28 (33.0) | 4 | 4 | 7 | 7 | 3 | 2 | 1 | 0 |
| Bronchial | 9 (10.6) | 0 | 2 | 3 | 4 | 0 | 0 | 0 | 0 |
| Mesenteric | 6 (7.0) | 1 | 2 | 1 | 2 | 0 | 0 | 0 | 0 |
| Lung | 16 (18.8) | 3 | 3 | 4 | 6 | 0 | 0 | 0 | 0 |
| Total | 85 (100.0) | 11 | 17 | 22 | 30 | 3 | 3 | 2 | 1 |

**Figure 2:**
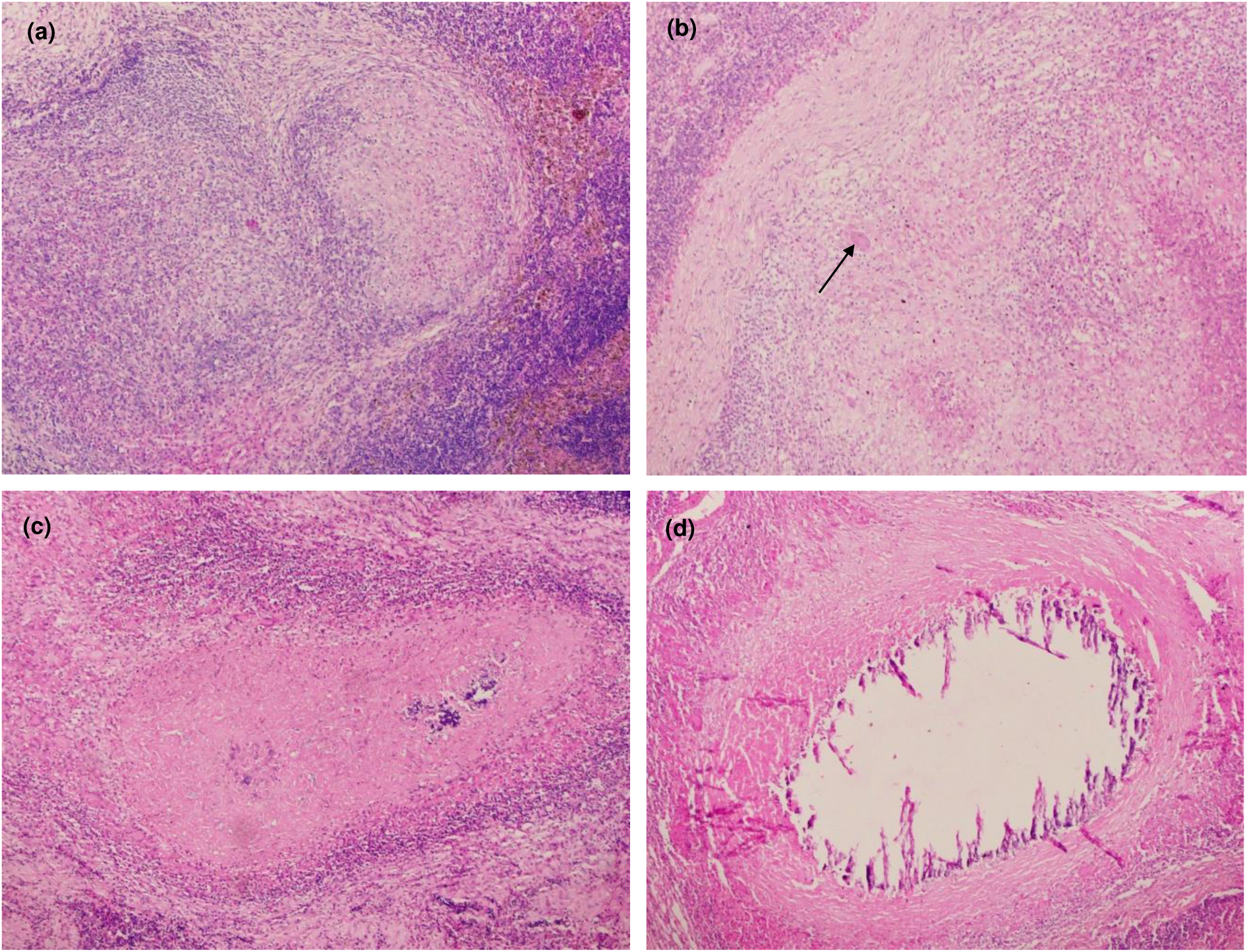
The four stages of granulomas in lymph nodes from naturally infected asymptomatic cows with *M. bovis*. a) Stage I (Initial). Clustered epithelioid macrophages are typical of this stage. HE 10*10. b) Stage II (Solid). Increased number of epitheliod macrophages including Langharn’s giant cells (arrow). Encapsulation is complete and central caseous necrosis is lacking. HE 10*10. c) Stage III (Minimal necrosis) thinly encapsulated with epitheliod macrophages and caseous necrosis. HE 10*10. d) Stage IV (Necrosis and mineralization). Large, irregular, encapsulated granuloma, often with multiple centers of caseuous necrosis and mineralization. HE 10*10.

### Acid fast bacillus staining

The presence of acid-fast bacilli in the tissue was examined by modified Zeihl Nelseen technique and there was no correlation between the stage of the granuloma and the AFB positivity (Fig. 3).

**Figure 3:**
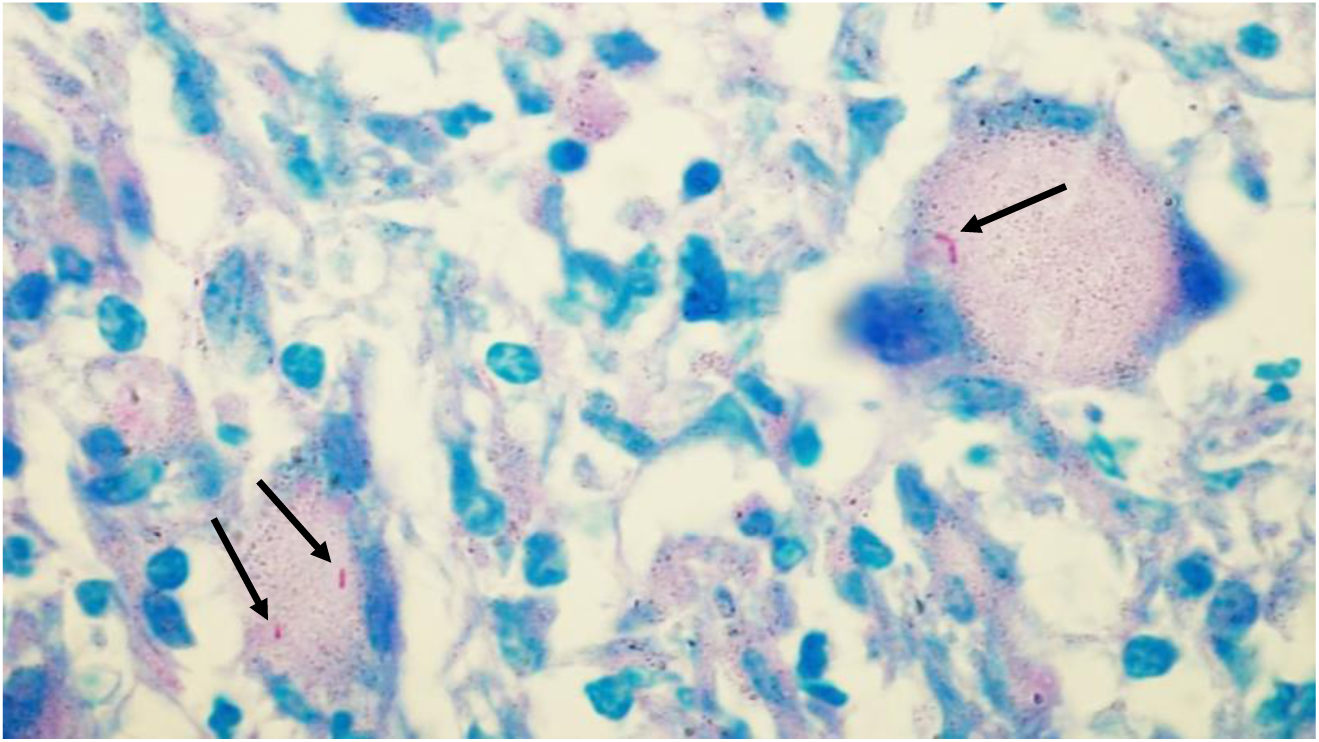
Acid fast bacilli (AFB) accumulate in the granuloma of tissue obtained from cows naturally infected with *M. bovis*. Intracellular, red colored bacilli indicated by black arrow.

### Immunohistochemistry

In this section, the cellular profiles and cytokine expressions were investigated in the granulomatous lesions in lymph nodes and lung of cattle naturally infected with *M. bovis*. The expressions of these cytokines and cellular profiles were evaluated in different stages of granuloma as well as in granulomas of culture positive and culture negative animals. Accordingly, CD3+ T cells and CD68+ macrophages and interferon gamma (IFN-γ), tumor necrosis factor alpha (TNF-α) and inducible nitric oxide synthase (iNOS) were evaluated (Table 2). The negative controls did not show positive labeling.

**Table 2:** Median percentage of immunolabeling of different cellular, cytokine and effector molecule in the granulomas (stage I to IV) of naturally infected bTB culture positive and negative cows. Data are presented as median ± SD

| Stages of granuloma | Median ± SD of immunolabeling | | | | |
| --- | --- | --- | --- | --- | --- |
| | CD3+ | CD68+ | IFN-γ+ | TNF- $\alpha$+ | iNOS+ |
| Stage I |  |  |  |  |  |
| Culture positive | 26.48 ± 4.1 | 4.21 ± 1.4 | 2.98 ± 1.1 | 17.73 ± 8.1 | 26.39 ± 9.9 |
| Culture negative | 23.12 ± 7.8 | 5.58 ± 2.6 | 3.77 ± 0.8 | 17.99 ± 9.2 | 41.89 ± 10.1 |
| Stage II |  |  |  |  |  |
| Culture positive | 21.5 ± 3.4 | 4.96 ± 1.6 | 8.81 ± 4.9 | 28.87 ± 6.2 | 44.96 ± 6.8 |
| Culture negative | 19.18 ± 4.6 | 7.56 ± 1.5 | 12.5 ± 5.4 | 30.85 ± 10.1 | 47.81 ± 10.2 |
| Stage III |  |  |  |  |  |
| Culture positive | 14.8 ± 4.0 | 6.69 ± 1.7 | 30.79 ± 7.7 | 25.93 ± 6.0 | 46.22 ± 3.7 |
| Culture negative | 19.76 ± 3.9 | 9.04 ± 1.7 | 36.12 ± 8.1 | 35.35 ± 8.4 | 46.54 ± 5.4 |
| Stage IV |  |  |  |  |  |
| Culture positive | 11.34 ± 2.4 | 9.77 ± 2.4 | 36.49 ± 6.5 | 32.05 ± 7.4 | 50.67 ± 6.5 |
| Culture negative | 19.07 ± 4.0 | 10.39 ± 4.1 | 42.88 ± 9.2 | 30.28 ± 8.2 | 43.69 ± 9.1 |

### T cells (CD3+)

Based on one way ANNOVA, in culture positive animals the mean CD3+ proportion fraction was seen to decrease as the stage of granuloma increased from stage I to stage IV (p<0.001) (Fig. 4). In culture negative animals, the mean difference of CD3+ proportion fraction as the stage of granuloma increase from stage I to IV, was not statistically significant (p>0.05). On the other hand, it was noted that at the advanced stage granulomas, there was a higher proportion of CD3+ cells in culture negative cows compared to culture positive animals (p<0.001).

**Figure 4:**
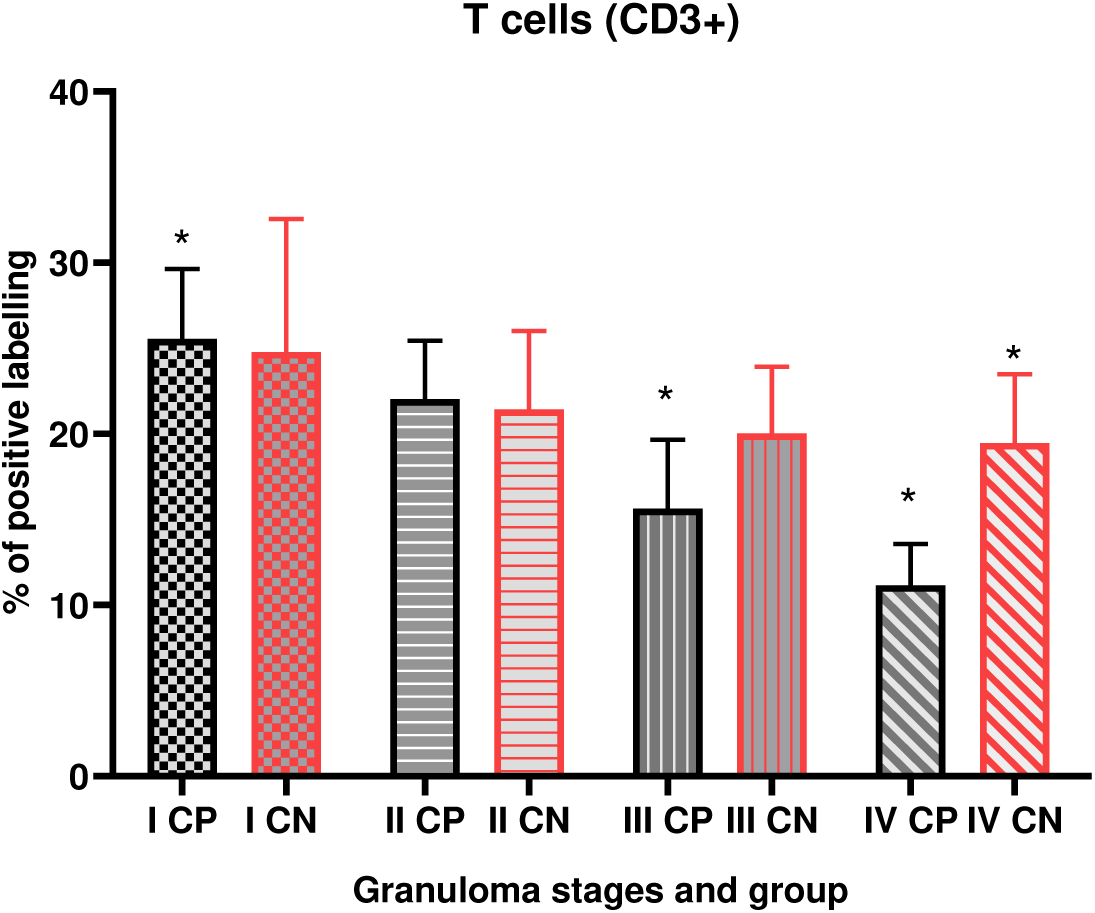
T cells (CD3+). Mean percentage of immunolabeling with in granulomas of stage I to IV for CD3+ T cells within the lymph nodes and lung tissue. The results are expressed as mean and SD. * p<0.05.

### Macrophages (CD68+)

Immunohistochemistry labelling of CD68+ identifies the locations of epithelioid macrophages and multinucleated giant cells (MNGCs). As the level of granuloma increases from stage I to IV, the proportion fraction of CD68+ immulobaleling increases with in the granuloma of both culture positive and negative animals (Fig. 5). The result obtained was statistically significant for culture positive animals (p<0.001) but not for culture negative animals (p>0.05). On the other hand, at the stage I granuloma, we observed a statistically significant (p=0.033) mean difference by comparing CD68+ immunolabling between granulomas from culture positive and culturing negative cows.

**Figure 5:**
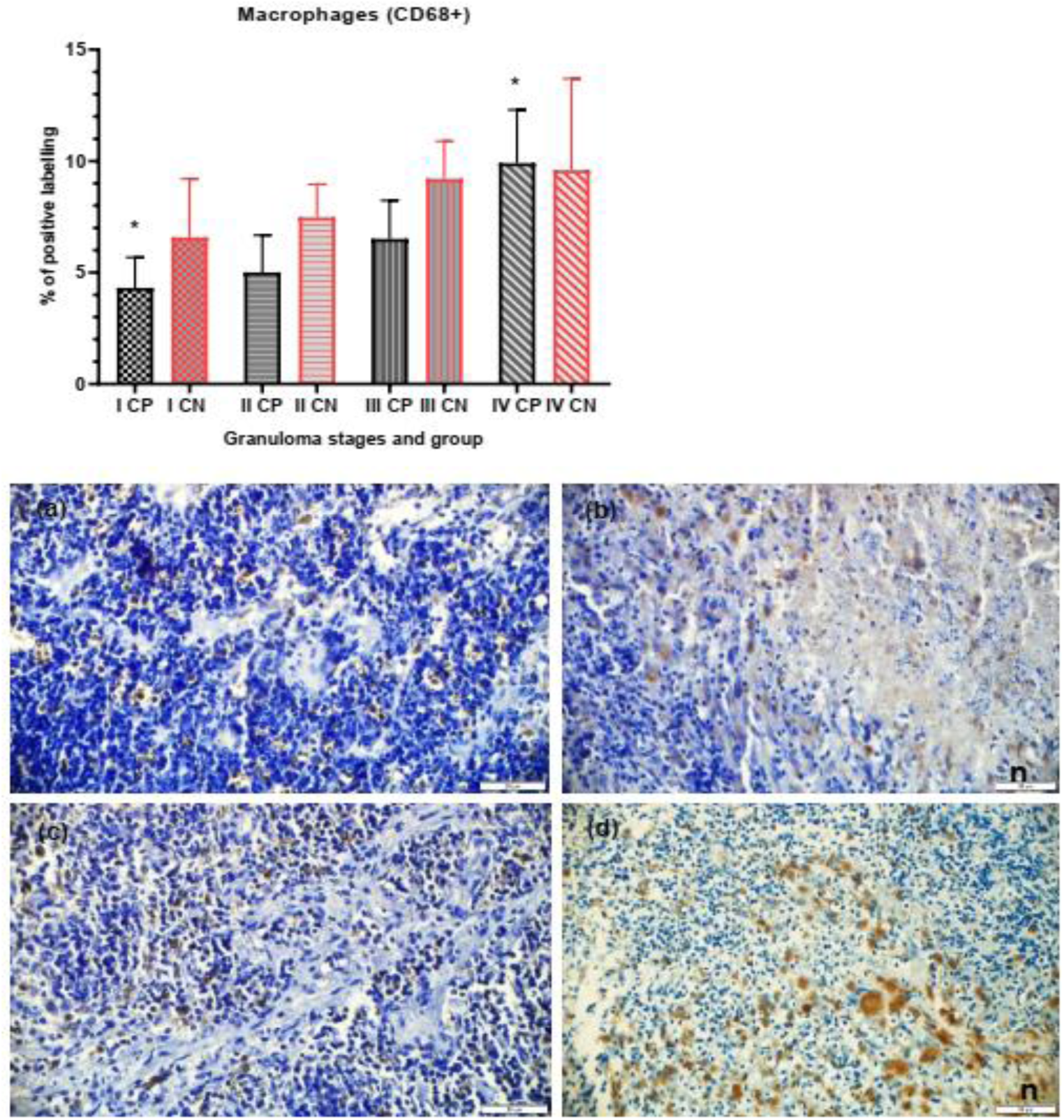
Macrophages (CD68+). Mean percentage of immunolabeling with in granulomas of stage I to IV for CD68+ with in the lymph nodes and lung tissue. The results are expressed as means and SD. *p<0.05. Immunlabeling of CD68+ macrophages of the lymph nodes of *M. bovis* culture positive (a, b) and (c, d) culture negative animals. Higher percentages of CD68+ macrophages can be seen in stage IV granulomas (b, d) compared to stage I (a, c).

### Cytokines IFN-γ+ and TNF-α+

Immunohistochemical labelling for IFN-γ+ (Fig. 6) showed increased mean percentage of positive labelling for granulomatous lesions from stage I to IV for both culture positive and culture negative cows (p<0.001). The IFN-γ+ mean difference between culture positive and culture negative animals was not statistically significant (p>0.05). With regard to TNF-α+ (Fig. 7), for culture positive cows, a statistically significant increase in mean percentage of immunolabelling was observed as the granuloma increase from stage I to IV (p< 0.001). While for culture negative cows, the mean percentage of TNF-α+ was not statistically significant as the level of granuloma increases.

**Figure 6:**
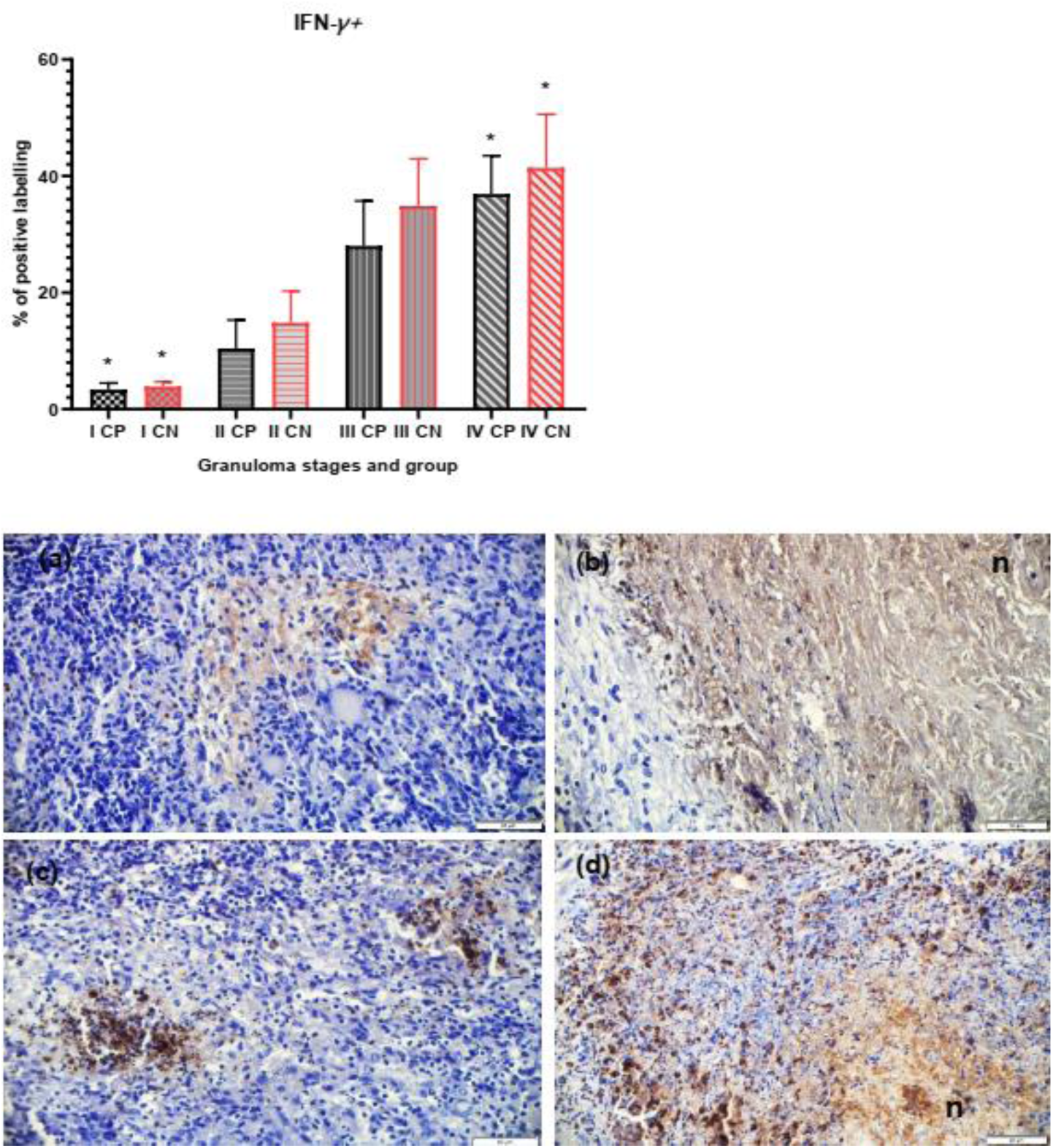
Interferon gamma (IFN-γ+). Mean percentage of immunolabeling with in granulomas of stage I to IV for IFN-γ+ with in the lymph nodes and lung tissue. The results are expressed as means and SD.* p<0.05. Immunolabeling of IFN-γ+ cells of the lymph nodes of *M. bovis* culture positive (a, b) and (c, d) culture negative animals. Higher percentages of IFN-γ+ cells can be seen in stage IV granulomas (b, d) compared to stage I (a, c).

**Figure 7:**
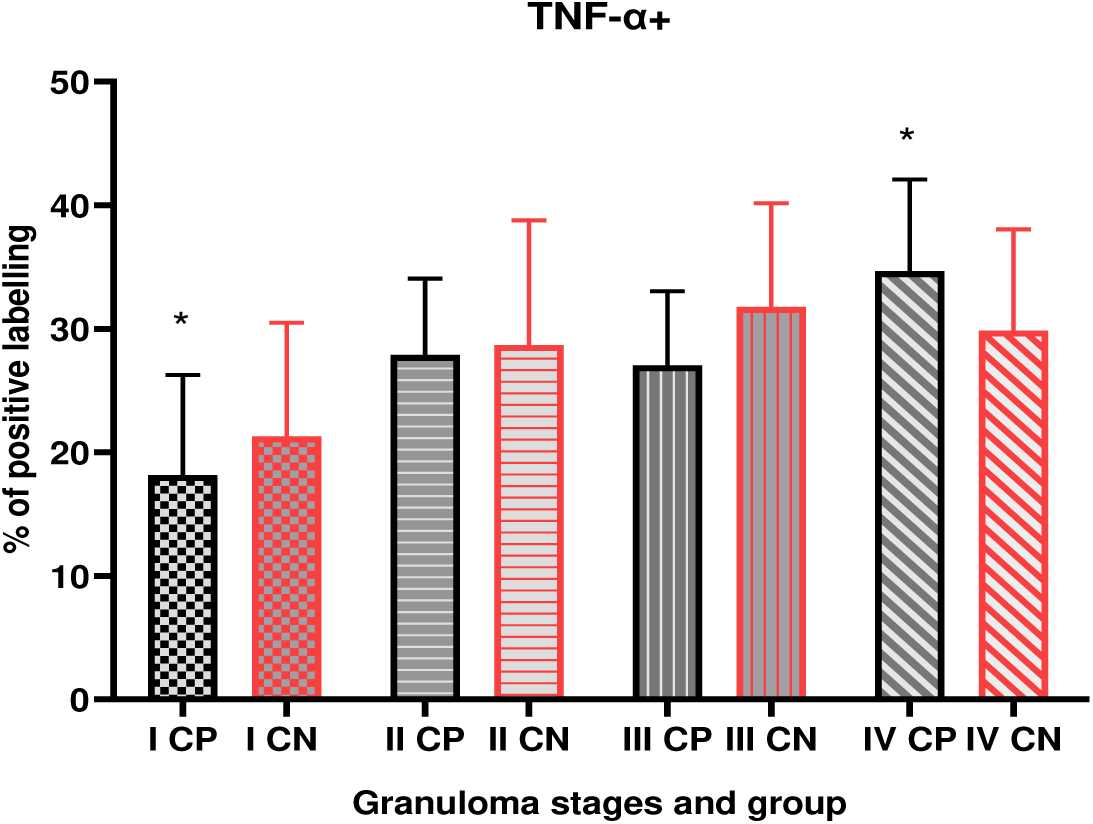
Tumor necrosis factor-alpha (TNF-α+). Mean percentage of immunolabeling with in granulomas of stage I to IV for TNF-α+ with in the lymph nodes and lung tissue. The results are expressed as means and SD.

### Inducible nitric oxide synthase (iNOS+)

Immunohistochemical staining of iNOS (Fig. 8) showed that the mean percentage of the labeling increases with statistically significant for culture positive animals from stage I through stage IV (p=0.0001). However, culture negative animals did not show significant difference in mean percentage of iNOS+ labeling as the stage of granuloma increases from stage I to stage IV.

**Figure 8:**
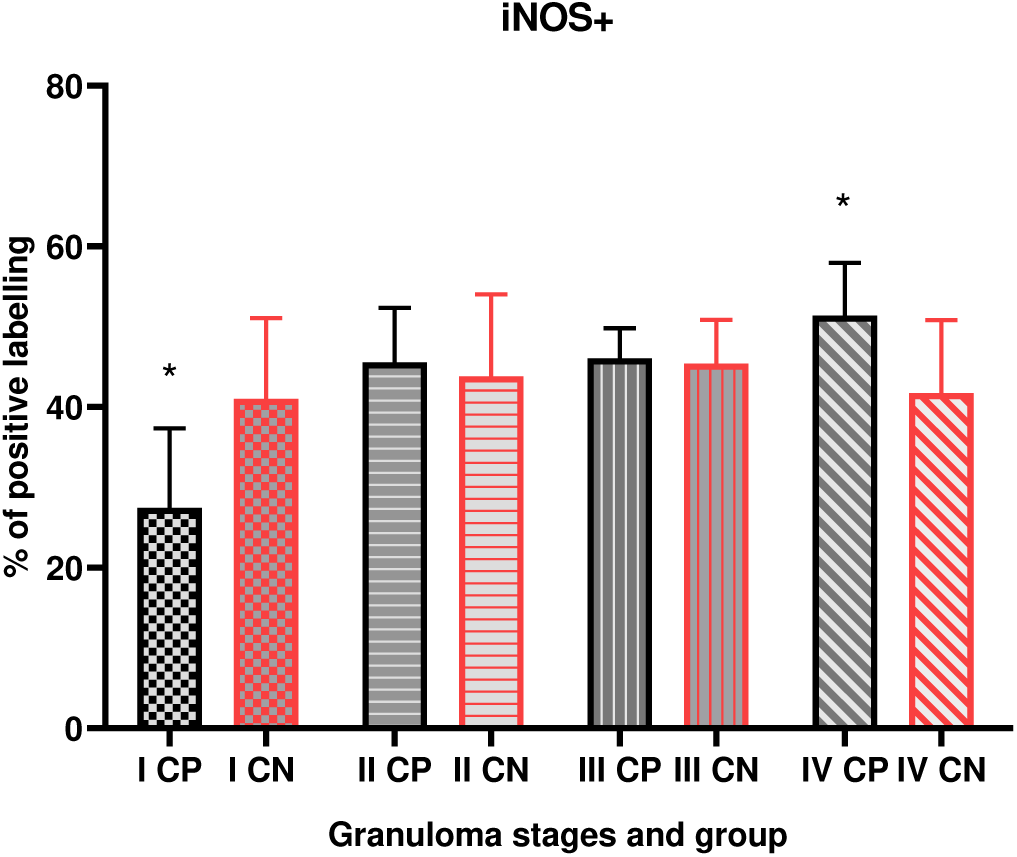
Inducible nitric oxide synthase (iNOS+). Mean percentage of immunolabeling with in granulomas of stage I to IV for iNOs+ with in the lymph nodes and lung tissue. The results are expressed as means and SD. *p<0.05.

## DISCUSSION

Evaluations of the lymph node using gross pathology, histological scoring and immunohistochemical markers during bTB lesions are useful to assess the immune response and classify granulomas to understand the diseases progress and severity (36). Moreover, it is important to determine vaccine efficacy and developments of immune-dependent diagnostic tools (19). Immunohistochemistry is one of the most commonly used tools to investigate cellular and molecular components of granuloma bTB lesion developed under experimental setup. However, information related to cellular and molecular aspect of the host immune response during natural infection is few. Studies showed that immune response of cattle against *M, bovis* natural infection may be affected by several host factors (33, 36).

In both culture positive and culture negative groups of animals, TB lesions were found to be more frequent and severe in the thoracic lymph nodes, supporting that respiratory route is the most common route of infection (31). More specifically, the caudal mediastinal, bronchial and cranial mediastinal lymph nodes displayed higher gross and microscopic scores in frequency gross pathology scoring and microscopic severity. This result is in agreement with study reported from similar study area by Ameni et al., (2006) where naturally *M. bovis*-infected cattle exposed to intensive husbandry system demonstrated higher frequency and severity of bTB-lesions in the respiratory tract. On the other hand, cattle kept on pasture showed higher severity of bTB lesions in their abdominal lymph nodes (33, 37). In this study the involvement of head and abdominal lymph node is relatively higher in culture positive group than culture negative group, suggesting the potential role of oral and other infection routes.

In immunohistochemical examinations it was observed that in culture positive animals, the immunolabeling fraction of CD68+ macrophages increased as the level of granuloma increases from stage I to stage IV in *M. bovis*. However, in *M. bovis* culture negative animals, it was noted that the pattern of CD68+ population remains the same across the four granuloma stages. A similar result was reported by Wangoo et al. (2005) where their experimental infection observed that CD68+ cells increase in number as the level of granuloma increases (27). The presence of large number of MNGCs in advanced granuloma in culture positive groups compared to culture negative group could be an indication of the active multiplication of the *M. bovis* bacteria when the immune response is not able to contain the microorganism (38).

In contrast to CD68+ macrophages, the proportion of CD3+ cells decreases as the stage of granuloma increases from stage I to stage IV; highest in stage I and lowest in stage IV in culture positive cows. However, in *M. bovis* culture negative granulomas, it was noted that there was no significant decrease in the mean population of CD3+ T cells population surrounding the granuloma as the stage of granuloma increases. This finding is supported by an experimental study designed to evaluate the role of CD3+ cells response in BCG vaccinated and non-vaccinated groups during *M. bovis* infection (28), suggesting the role of adaptive immune response mediated by T cells in containment of *M. bovis* infection. Most importantly, the cell-mediated immune response effected by CD4+ T cells by producing Th1 cytokines, such as IFN-γ, and the cytolytic activity of CD8+ cells toward infected macrophages is crucial (39).

Immunohistochemical examinations of IFN-γ+, TNF-α+ and iNOS+ shows the same immunolabeling pattern as that of CD68+ macrophages, which increases as the level of granuloma increases from stage I to stage IV in culture positive animals. Evidence from natural *M. bovis* infection from other species has shown that the presence of CD68 macrophages and CD3 T cells in and surrounding granuloma is in agreement with the high level expression of pro-inflammatory cytokines like IFN-γ and TNF-α and iNOS effector molecules (34). These pro-inflammatory cytokines are important in promoting the formation and function of granuloma. Previous studies (27, 28, 35) observed a significant increase in the level of pro-inflammatory cytokine, mainly IFN-γ+, as the stage of granuloma advances. Also nitric oxide (NO) production by macrophages during mycobacterial infection play a crucial role in the intracellular killing of mycobacteria as it is cytotoxic at high concentrations (23). This observed increase in pro-inflammatory cytokines (IFN-γ and TNF-α) and effector molecules (iNOS) seems likely to have contributed to the regulation bovine immune response during *M. bovis* infection (35).

Evidence from this study provides basic information on the host immune response during natural infection with *M. bovis*, which could be used for further studies in the investigation of biomarkers necessary for diagnostics and vaccines in the fight against bTB. However, this study may not be generalized to other countries but could reflect the situation in Ethiopia. The small number of culture negative animals used as comparison may also limit the generalizability of this study.

## CONCLUSION

This study highlighted the role of macrophages, T cells and chemical mediators like IFN-γ, TNF-α and iNOS during naturally infected asymptomatic cows with *M. bovis* from intensive dairy farms in central Ethiopia. For *M. bovis* culture positive animals, the activity of CD68 macrophages, IFN-γ, TNF-α and iNOS were more intense as the level of granuloma increases while CD3 T cells population decreases as the stage of granuloma increases. Thus, the activity of CD68+, IFN-γ+, TNF-α+ and iNOS+ could play a protective role in the immune defense against *M. bovis* during naturally infected asymptomatic cows.

## MATERIAL AND METHODS

### Study setting and ethical statement

The study was conducted on semi-urban intensive dairy farms situated in central Ethiopia, Oromia Special Zone surrounding Addis Ababa City, the capital of Ethiopia. The study obtained ethical approved from the Armauer Hansen Research Institute (AHRI) Ethics Review Committee (Ref P018/17), from the Ethiopian National Research Ethics Review Committee (Ref 310/253/2017), the Queen Mary University of London Research Ethics Committee, London UK (Ref 16/YH/0410); and by the Aklilu Lemma Institute of Pathobiology, Addis Ababa University (Ref ALIPB/IRB/011/2017/18). Written informed consent was obtained from all the owners of the farms.

### Animals

A total of 16 single intradermal cervical comparative tuberculin test (SICCTT) positive cows suspected to be naturally infected with *M. bovis* were obtained from 16 different farms. Sex and body condition score (BCS) were recorded. A method developed by Nicholson and Butterworth (40) was used to determine the BCS. Poor BCS was considered with extremely lean cattle with projecting dorsal spines pointed to the touch and individual noticeable transverse processes. Medium BCS was considered with cattle with usually visible ribs having little fat cover and barely visible dorsal spines. Good BCS was considered with Fat cover is easily observed in critical areas and the transverse processes were not visible or felt.

### SICCTT

Briefly, SCITT was performed as follows. Two sites on the right side of the mid-neck, 12 cm apart, were shaved, and the skin thicknesses were measured with calipers. One site was injected with an aliquot of 0.1ml containing 2,500 IU/ml bovine PPD (PPDb) (Veterinary Laboratories Agency, Addlestone, Surrey, United Kingdom). Similarly, 0.1ml of 2,500 IU/ml avian PPD (PPDa) (Veterinary Laboratories Agency, Addlestone, Surry, United Kingdom) was injected into the second site. After 72 h, the skin thicknesses at the injection sites were measured (11). Then the difference between the swellings of PPDa and PPDb were calculated and the positive result was determined at cut off ≥ 2 mm.

### Postmortem examination

The cows were humanely slaughtered by personnel of the local abattoirs in the study area. The post-mortem examination was performed by an experienced meat inspector. From all the 16 animals, a total of a total of 176 lymph nodes and 96 lung tissues were examined by slicing the tissue into 0.5-1cm sections, and assigning a pathology severity score, as developed by Vordermeier *et al*., 2002 (30) shown in Table S2. Both lymph node and lung pathology score were added to determine the total pathology score per animal. In order to maintain the scoring consistency, all scoring was performed by a single person.

### Culture

Isolation of mycobacteria was performed according to World Organization for Animal Health protocols (41). Briefly, tissue specimens for culture were collected into sterile universal bottles in 5ml of 0.9 % saline solution, and then transported to Aklilu Lemma Institute of Pathobiology (ALIPB) TB laboratory. The tissues were sectioned, homogenized and the sediment was neutralized by 1% (0.1N) HCl using phenol red as an indicator. Thereafter, 0.1ml of suspension from each sample was spread onto a slant of two Löwenstein Jensen (42) medium tubes one enriched with sodium pyruvate and the other enriched with glycerol. Cultures were incubated aerobically at 37°C for at least eight weeks and with weekly observation of the growth of colonies. In order to report culture negative, the tissues were repeatedly cultured three times.

### Histopathology

A total of 37 tissue samples (27 culture positive and 10 culture negative) with high gross pathology scores were selected from lymph nodes and lung tissues. Lesions were carefully selected to include the encapsulated granulomas of different sizes with caseous necrosis and calcification.

The tissues were fixed in 10% neutral buffered formalin for 24-72 hours, embedded in paraffin, sectioned in 4μm sections and stained with hematoxylin-eosin (H&E) and Ziehl Neelsen acid fast stain. Granulomas were classified into different stage I to IV according to the previously described criteria (27). Acid fast bacilli (AFB) were recorded as being present or not.

### Immunohistochemistry

For the immunohistochemistry experiment, 4 μm formalin fixed tissue samples were stained with avidin-biotin-complex (ABC Vector Elite; Vector Laboratories) method. Tissue sections were first either deparaffinized or dewaxed and rehydrated. Antigen retrieval was induced by heat (Microwave) or enzymes (trypsin /chymotrypsin) (Sigma, Poole, UK) (Table S3) and adjusted to pH 9 or 6 using 0.1N sodium hydroxide. Tissue sections were washed in running tap water, and then incubated with a blocking buffer (normal goat/horse serum in 10 ml PBS) for 30 minutes. Slides were incubated with primary antibody for overnight and secondary antibody for 20 minutes and then the staining was amplified using avidin-biotin-peroxidase conjugate (ABC elite; Vector Laboratories) and visualized using 3, 30-diaminobenzidine tetrahydrochloride. The unbound conjugates were removed prior to DAB application with two buffer washes. Finally, the slides were washed in tap water and stained by Mayer’s Haematoxylin counterstain, and mounted for analysis. For negative control tissue we used a bovine lymph node with no gross lesion and no isolation of *M. bovis* with culture. For each experiment we included a slide with secondary antibody but no primary antibody.

### Image analysis

For each granuloma, a total of 10 fields from different areas of the granuloma, avoiding necrotic and mineralized areas, were analysed using a Fiji-ImageJ software (https://imagej.net/Fiji/Downloads). All images were examined at X400 magnification, and captured with an Olympus®DP74 digital camera attached to a microscope BX Olympus®63. Briefly, after image was imported to Fijii-Image J software actual color was deconvulated into three different colors (green, gray and blue) using H DAB vector. The second color (gray) used for further processing and converted into black and white contrast using “Make Binary” tool, color threshold was adjusted at default (0 scale for min and 255 for max). Next mean (including minimum and maximum) value of area of fraction was taken and percent area was determined (43). For each antibody, the total area of positive labelling was given as a percentage of the total area examined in 10 fields.

### Statistical Analysis

The results of the histopathological and the immunohistochemical analysis were expressed in mean and standard deviation, and the results were compared between the stages of granuloma and between culture results. A nonparametric statistical analysis employing One-Way ANOVA and unpaired t-test were used to compare the means and p<0.05 was considered statistically significant. The analyses were conducted using GraphPad Prism 8.0 (San Diego, CA, USA).

## CONTRIBUTORS

BT and GA conceived the study. BT, AZ, FD, HMM, ARM and GA contributed to study design and development of laboratory assays. BT, AZ, FD, HMM, MB, MA, TT, DAG, ARM and GA contributed to implementation of the study and contributed to the data acquisition. BT did statistical analyses, wrote the first draft of the manuscript and had final responsibility for the decision to submit for publication. All authors reviewed the final draft and agree with its content and conclusions.

## ACKNOWLEDGMENTS

This work was supported by a grant from the United Kingdom Medical Research Council (Reference Number MR/P024548/1, to ARM). We thank all the members of the field and laboratory teams at AHRI and ALIPB; Mr Mengistu Mulu, Lemma Terfasa and Mr Tadesse Regassa for assistance with field work and postmortem examination; Mrs Yemisrach Zeleke for assistance with *M. bovis* culture; Mrs Sofia Yimam and Mr Selfu Girma for assistance with histopathology, AFB examination and immunohistochemistry.

